# Deep learning-based Spatial Feature Extraction for Prognostic Prediction of Hepatocellular Carcinoma from Pathological Images

**DOI:** 10.1101/2024.02.10.579571

**Authors:** Huijuan Hu, Tianhua Tan, Yerong Liu, Wei Liang, Wei Zhang, Ju Cui, Jinghai Song, Xuefei Li

**Author notes:** These authors contributed equally (Huijuan Hu, Tianhua Tan).

## Abstract

The spatial structures of various cell types in tumor tissues have been demonstrated to be able to provide useful information for the evaluation of the disease progression as well as the responsiveness to targeted therapies. Therefore, powered by machine-learning, several image segmentation methods have been developed to identify tumor-cells, stromal, lymphocytes, etc., in hematoxylin and eosin (H&E) stained pathological images. However, the quantitative and systematic characterization of the spatial structures of various cell types is still challenging. In this work, we first developed a robust procedure based on deep learning to precisely recognize cancer cells, stromal and lymphocytes in H&E-stained pathological images of hepatocellular carcinoma (HCC). In order to quantitatively characterize the composition and spatial arrangement of the tumor microenvironment, we then systematically constructed 109 spatial features based on locations of the 3 major types of cells in the H&E images. Interestingly, we discovered that the absolute values of several spatial features are significantly associated with patient overall survival in two independent patient cohorts, such as the cellular diversity around stromal cells (StrDiv), the average distance between stromal cells (StrDis), the coefficient of variation of the tumor-cell polygon area in the Voronoi diagram (TumCV), *etc*., based on univariate analysis. In addition, multivariate Cox regress analyses further demonstrated that StrDiv and StrDis are independent survival prognostic factors for HCC patient from The Cancer Genome Atlas Program (TCGA). Furthermore, we demonstrated that a combination analysis with cell spatial features, *i.e*. StrDiv or TumCV, and another important clinical feature, *i.e*. microvascular invasion (MVI), can further improve the efficacy of prognostic stratification for patients from the Beijing Hospital cohorts. In summary, the spatial features of tumor microenvironment enabled by the digital image analysis pipeline developed in this work can be effective in patient stratification, which holds the promise for its usage in predicting the therapeutic response of patients in the future.

## Introduction

Hepatocellular carcinoma (HCC) is the most prevalent form of primary liver cancer, accounting for 75% to 85% of cases worldwide, and it ranks as the sixth most common malignancy and the fourth leading cause of cancer-related mortality [1]. HCC exhibits significant heterogeneity at the histological, molecular, and genetic levels, posing challenges for accurate prognostic stratification and personalized therapy. Consequently, there is a growing emphasis on exploring new methods for more accurately stratifying cancer patients into distinct prognostic groups with different predicted outcomes [4]. Previous attempts to stratify HCC patients based on genomic, transcriptomic, and proteomic profiles have yet to fully unveil the links between these molecular traits and appropriate therapeutic interventions [2, 3].

Other than molecular traits, histopathology images have been widely employed for cancer diagnosis and prognosis, since it can provide morphological features of cells that are highly related to the degree of the aggressiveness of cancers [5,6]. Moreover, these images can offer spatial-structure information about the tumor microenvironment as the locations of individual cells are available [7-9]. The interaction between tumor cells and their microenvironment plays a pivotal role in cancer development and progression [10, 11]. Recent studies have demonstrated the diagnostic and prognostic value of lymphocytes and their spatial distribution in various cancers [10-15]. Additionally, the spatial distribution pattern of stromal cells in pathological images has shown effectiveness in predicting patient prognosis [16,17].

Despite the wealth information contained within pathological images, manually labeling single cells and extracting quantitative spatial features of the tumor microenvironment is infeasible. In recent years, Deep learning algorithms have emerged as a promising solution to address this challenge. Specifically, deep learning-based approach has been applied to simultaneously recognize and segment tumor cells, lymphocytes, and other non-malignant cells in regular pathological images of HCC [22]. However, the recognition accuracy of non-malignant within the tumor microenvironment is relatively low [22]. More importantly, quantitatively characterizing the complex tumor microenvironment based on the spatial coordinates of multiple cell types remains a challenge. Addressing these limitations and advancing the field towards more precise and comprehensive characterization of the tumor microenvironment is a critical area of cancer research [23, 24].

This study aims to address this challenge by simultaneously extracting spatial information of multiple cell types in regular pathological images of HCC patients, and quantitatively and systematically characterizing the tumor microenvironment based on cell locations. By utilizing image processing techniques [25] and modifying a cell recognition convolutional neural network (CNN) [29] initially designed for lung cancer, we have created a computational pipeline that can precisely detect lymphocytes, stromal cells, and tumor cells from H&E tumor section images of HCC patients.

Moreover, by analyzing the spatial characteristics of the three cell types with the help of Delaunay network and Voronoi diagram, we have successfully constructed 109 spatial features. Among them, cellular diversity around cells in the Delaunay network (CellDiv) and coefficient of variation of the cell polygon area in the Voronoi diagram (CellCV) represent spatial heterogeneity in the hypercellular network. Specifically, we showed that the higher the cellular diversity around stromal cells (StrDiv), the better the patient’s prognosis is. In addition, the lower the coefficient of variation of the tumor-cell polygon area in the Voronoi diagram (TumCV), i.e., the more homogeneous the tumor cells distribute, the longer the patient’s overall survival is. This was validated in our univariate Cox regression analysis for both TCGA HCC cohorts and Beijing Hospital one. Furthermore, multivariate regression analysis for both cohorts further demonstrate that StrDiv could be an independent factor for predicting patient overall survival. In addition, continuous variable regression also indicates that these variables are also significantly correlated with patient overall survival. Last but not least, when combining with other important clinical prognostic factors, such as microvascular invasion (MVI), TumCV and StrDiv can help us to further increase the precision of stratification for patients with and without MVI, respectively.

In summary, our study presents a novel computational pipeline capable of digitalizing regular H&E tumor-section images of HCC patients and extracting quantitative human-interpretable spatial features of the tumor microenvironment. These findings hold the potential to develop diverse prognostic biomarkers based on digital pathology.

## METHODS

### Patient information

There are two cohorts of HCC patients in this study. The first cohort involved 207 patients from publica data base (TCGA) under the category of LIHC, and their patient ID were listed in Supplementary Table S1. Corresponding Formalin-fixed paraffin-embedded (FFPE) whole-tumor slide images (WSIs) were downloaded from https://www.cancer.gov/tcga. The second cohort involved 69 HCC patients with whole-tumor pathological slide images. All patients received curative hepatectomy between 2012 and 2020 at Beijing Hospital, and they did not have distant metastasis or any prior anticancer treatments before the surgery. All the images were FFPE slides and digitally captured at 40X magnification. As for the clinical variables are preprocessed in the following way. Pathologic stage was mapped into three groups as well: early (stage I, Ia, Ib), locally advanced (stage II, IIa, IIb) and advanced (stage IIIa, IIIb, IV). Others was shown in Supplementary Table S4.

### Histopathologic image-Processing Pipeline

The ImageScope annotation tool was used to manually label boundaries of region of interest (ROI). ROIs were defined by the major malignant region in each pathological image. Within each ROI, we then randomly sampled 20 patches, whose dimension were 5000×5000 pixels under 40X magnification (0.25 μm per pixel).

To obtain the location of each cell nucleus, we first adopted a deconvolution method [25] to convert the RGB colors of the H&E image into the Hematoxylin and Eosin (HE) color space, with the deconvolution matrix [0.6500, 0.7040,0.2860; 0.2681, 0.5703, 0.7764; 0.7110, 0.4232, 0.5616], where signals of cell nucleus would be enriched in the Hematoxylin channel. Then, to reduce the fragments of cell nucleus caused by noise, morphological operations consisting of opening and closing [26] were used to process the hematoxylin channel image. Subsequently, FastMarching segmentation method [27] was applied to detect nuclei boundaries. The parameters of the segmentation algorithm are typically set as follows: sigma=0.5, alpha=-0.3, beta=2.0, threshold=200, stopping time=210. Seed position was the coordinate of any point in the expected segmentation area. Finally, Image patches with the size 80×80 centered on the detected nuclei center were extracted from the original selected pathological RGB image (illustrated in Fig. 1). Such image-processing procedure can improve the accuracy of cell recognition by the deep-learning procedure.

**Fig. 1.**
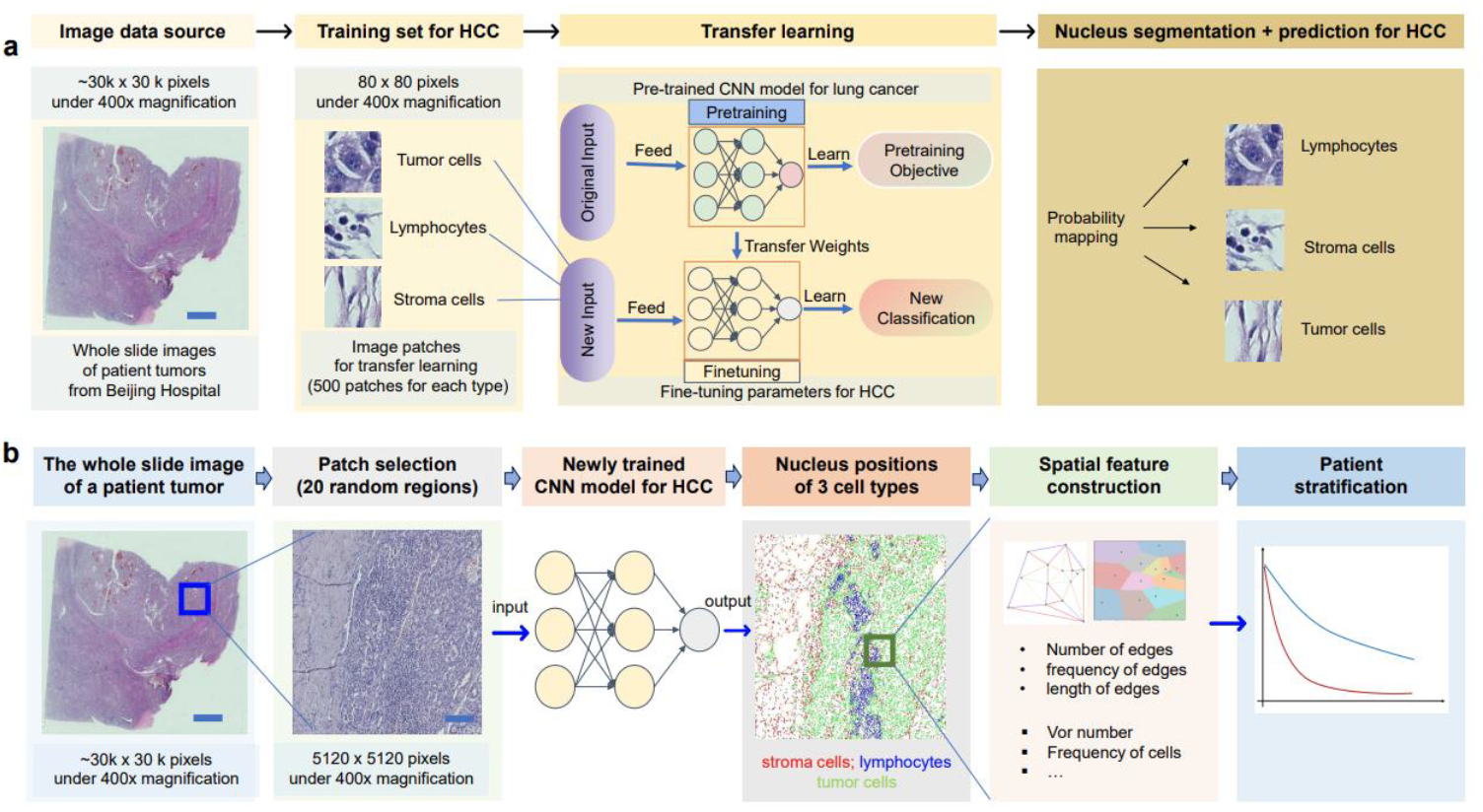
Data and workflow for the prognostic analysis of HCC. (a) The cell classification method includes nuclei identification through HD-Staining, Transfer-learning, and classification. (b) Randomly select 20 regions of ROI to segment and classify each nucleus by HCC-CNN model. Construct Delaunay triangulation and Voronoi diagram to obtain 109 quantitative spatial features. Then explore the correlation of proposed features with the prognosis of cancer patients.

### Transfer learning CNN model

To effectively and simultaneously recognize lymphocytes, tumor and stromal cells in H&E images of HCC patients, we implemented a Transfer Learning CNN model, which was developed for recognizing cells in lung cancer [28]. All networks received an input image patch sized 80×80 which was normalized to the range [-0.5,0.5] with R, G, B channels. Same as the original CNN model, an 8-layer deep convolution neural network was applied which consists of three convolution layer, three pooling layers, one fully connected layer, and an output layer, as shown in Fig. 1. The pooling layer uses the maximum pooling and the output layer was a softmax layer with 3 categories: tumor cell, stromal cell, and lymphocyte. For each image patch, probability for each of the 3 categories was predicted and the highest probability was considered as the predicted class. During training, data augmentation including flip, rotation, Gaussian blur was applied to augment sample size. we initialized the model with pre-trained weights [28] on the lung adenocarcinoma datasets, and then fine-tuned parameters for softmax layer based on 500 patches of our marked examples on HCC datasets by Transfer learning [29].

### Construction of the topological features based on tumor-cell, stromal cell and lymphocyte

With the centroid of each cell-nucleus obtained, the triangulation algorithm Delaunay [32] were adopted to construct the spatial topological network, in which each nucleus and its adjacent nuclei were connected by edges. Meanwhile, we also applied Voronoi diagram [39] which divides the space into many areas based on the nearest attributes of the objects. In such diagram, the distance from any point within a convex polygon’s enclosed region to the object point of that polygon is less than the distance to any other object point.

Based on the space location of graph nodes in graph measurements (i.e., cell positions), we further constructed spatial features that capture the spatial arrangement and structural characteristics of the tumor microenvironment. In total, we constructed 109 spatial features, including cellular diversity around cells (CellDiv) (16), coefficient of variation of the cell polygon area in the Voronoi diagram (CellCV) (4), cell-neighbor-compass(16), edge distance of Delaunay network(35), frequency of edges (6), absolute cell numbers (11), cell polygon area in the Voronoi diagram (18), and cell density (3). A full list of these spatial features was listed in Supplementary Table S2. Note that since we selected 20 patches from the WSI of each patient, the topological features of cells for each patient were calculated based on the corresponding values in all patches from the same WSI. In the following, we showed two important categories of spatial features in details, i.e., cellular diversity around cells (CellDiv), and coefficient of variation of the cell polygon area in the Voronoi diagram (CellCV).

To describe the degree of cellular diversity in the tumor microenvironment, we adopted the quantitative measures proposed by Tang et al. [20]:

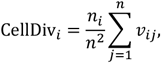

where *n*_*i*_ is the number of cell types in the *n* nearest neighboring linked cells of the target cell *i*, and *n* is the number of nearest neighboring linked cells of the target cell. When the target cell *i* and the nearest neighboring linked cell *j* belong to different cell types, *ν*_*ij*_ = 1, otherwise it is 0. A diversity heterogeneity of 0 (minimum value) indicates that there are no other types of cells as neighbors, while a diversity heterogeneity of 1 (maximum value) indicates that there is no same type of cells as neighbors. And we calculated the cellular diversity for each of the three cell types.

To describe the variance of the cellular influence, we adopted the measures proposed by DUYCKAERTS [40]:

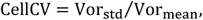

where Vor_std_ represents the standard deviation of Voronoi polygon areas and Vor_mean_ represents the average area of Voronoi polygons of a particular cell type. When the spatial distribution of cells and their neighbors is relatively consistent from one region to another, the area of the Voronoi polygons may have relatively low fluctuation and the CellCV value is relatively low, and vice versa. DUYCKAERTS[40] proposed 3 suggested values: when the point set is randomly distributed, the CellCV value is 33% to 64%, indicating random distribution; when the point set is clustered, the CellCV value is greater than 64%, indicating clustered distribution; when the CellCV value is less than 33%, the point set is uniformly distributed. And we also calculated the variance of cellular influence for each of the three cell types.

### Survival analysis

With the 109 spatial features calculated, we first separated each patient cohort into two groups with a specific cut-off value of a particular feature. Then the Kaplan-Meier (K-M) survival analysis and log rank tests were performed to test whether there exists a statistically significant difference in the overall survival (OS) between the two groups. The OS data were obtained from TCGA and Beijing Hospital, documented in Supplementary Table S1 and S5, and right censored. Specifically, for each feature, we selected the optimal cut-off value to divide the patients into two groups based on the method proposed by package survminerR. Specifically, the method uses the maximum selected rank statistic to test for independence between the response variable and a given feature. In addition, univariate and multivariate Cox proportional hazard model was employed to estimate the hazard ratios (HR) and 95 % confidence interval (CI). A two-side p value 0.05 was selected as the threshold for defining any statistical significance between patient groups. The survival risk ratio (hazard ratio, HR) was used to represent the difference in risk between different groups due to the variable intervention. All variables with log rank p < 0.05 on univariate analysis were entered for multivariate analysis with backward stepwise selection of variables. R (version 3.2.4) and R packages survival (version 2.38-3), glmnet (version 2.0-5), clinfun (version 1.0.13) and python (version3.5.2) were used in the analysis.

## Results

### Transfer-learning of the CNN model to accurately classify cells in H&E images of HCC

To accurately segment cell nucleus and recognize cell type using H&E-stained pathological images of HCC patients, we first adopted a transfer-learning procedure (see **Methods**), to recognize cancer cells, stromal, and lymphocytes in HCC (**Fig. 1**), based on a previously developed CNN model for tumor-section images of NSCLC [28]. Briefly, image patches were randomly selected from whole-slide H&E-stained pathological images of HCC patients at Beijing Hospital. Each patch is chosen based on the center around one specific type of cell nuclei, including lymphocytes, tumor, and stromal cells. In total, 500 patches for each cell type were selected as the training and validation sets for transfer-learning, and the cell type in each patch was confirmed by two pathologists at Beijing Hospital (**Fig. 1**).

For the transfer-learning procedure (see **Methods**), we adopted a cross-validation method, i.e., in each round of training and validation, 80% of the image patches were used as the training set, whereas the rest 20% were used as the validation set. As a result, the overall classification accuracy of the CNN model on training images were 96.3% for lymphocytes, 84.7% for stromal cells, and 89.1% for tumor cells, respectively. The independent cross-study classification rates in the test dataset were 92.3% for lymphocytes, 77.6% for stromal cells, and 95.7% for tumor cells.

### Spatial features of cells in tumor-section images of HCC patients

In total, we have two cohorts of patients, one is from Beijing Hospital including 67 patients, and the other one is from TCGA including 207 individuals (see **Methods**). For each patient, we collected one whole-slide image (WSI) of the primary tumor. For each WSI, we randomly selected 20 regions of interest (ROI), with each ROI having 5000 × 5000 pixels in size (under 400x magnification, see **Methods**). For each ROI, we applied our HCC-CNN model to segment and classify each cell nuclei and extracted the x-y coordinates of lymphocytes, tumor cells, and stromal cells. Based on these spatial coordinates, we used the Delaunay triangulation [32] as well as Voronoi diagram [39] to construct topological networks and diagrams of lymphocytes, tumor cells, and stromal cells (supp. 1(a-e)). Then, we created 109 quantitative image features to characterize the spatial distribution and relationships. A full list of these features and biological meanings were listed in Supplementary Table S2.

In addition to the simple features such as cell numbers, density, cell-cell distance, etc., there are a few unique features constructed. For example, we estimated the median value of overall cellular heterogeneity (MedMix), which describes the diversity of cell types around cells (ranging from 0-1, see **Methods**). The MedMix values for each of three cell types are 0.111 (tumor cells), 0.111 (stromal cells) and 0.083 (lymphocytes), which indicates that within the tumor microenvironment, the neighbors of each cell type tend to be cells of the same type. Moreover, with the Voronoi diagram, we were also able to calculate the Voronoi area of each cell nucleus (**Fig. 1b**) and thus the median and variance of such area for each cell type. Specifically, coefficient of variation of the cell polygon area in the Voronoi diagram for each of three cell types are 55.8% (tumor cells), 52.7% (stromal cells) and 21.6% (lymphocytes), indicating a strong fluctuation of cellular packing for tumor and stromal cells. In following sections, we will demonstrate that these unique spatial features can be important indicators of patient prognosis.

### Univariate analysis of the spatial features with patient prognosis on TCGA cohort

With the obtained spatial features of each patient, we explored whether these signatures could help to stratify patients into significantly different survival groups. We first conducted univariate analysis of the features using the Cox proportional hazards model (see **Methods**). Briefly, using the method proposed by package survminerR we can define the optimal cut-off value of each feature, which separates a patient cohort into two significantly different prognostic groups. We performed such analysis of all 109 spatial features for the 207 LIHC patients in TCGA and 67 patients for Beijing hospital. As a result, 48 out of 109 spatial features can effectively separate the 207 HCC patients from TCGA into two prognostic groups. For the Beijing hospital cohort, 70 out of the 109 spatial features can effectively separate the 67 patients into two prognostic groups. The detail list of the hazard ratio and p values can be found in Supplementary Table S3. Importantly, there are 13 spatial features that are identified for both cohorts of patients (Table 1). Note that the results shown in Table 1 were obtained with different optimal cut-off values of the corresponding features.

**Table 1.** Univariable analyses of factors associated with overall survival on TCGA cohort and Beijing cohort.

| Features | TCGA cohort |  |  | Beijing cohort |  |  |
| --- | --- | --- | --- | --- | --- | --- |
|  | CI | HR | P | CI | HR | P |
| Compass_mean | [1.01, 2.56] | 1.61 | 0.046 | [1.39,25.27] | 5.94 | 0.016 |
| llcompass_var | [1.08, 3.96] | 2.07 | 0.028 | [1.21,7.68] | 3.04 | 0.018 |
| LI_distance | [1.38, 3.58] | 2.22 | **<0.001** | [1.06,7.60] | 2.83 | 0.038 |
| ttcompass_var | [1.16, 3.23] | 1.94 | 0.011 | [1.57, 8.01] | 3.55 | 0.002 |
| sscompass_median | [1.06, 2.69] | 1.69 | 0.027 | [2.23,122.57] | 16.52 | 0.006 |
| StrDis | [1.15, 3.05] | 1.88 | 0.011 | [1.27, 6.68] | 2.92 | 0.011 |
| tumor_number | [0.33, 0.91] | 0.54 | 0.019 | [0.06, 0.47] | 0.17 | **<0.001** |
| tumor_density | [0.39, 0.99] | 0.62 | 0.044 | [0.11, 0.57] | 0.25 | **<0.001** |
| tumor_area | [0.36, 0.98] | 0.59 | 0.042 | [0.02, 0.87] | 0.12 | 0.036 |
| ll_frequency | [1.30, 3.43] | 2.11 | 0.002 | [1.09, 6.93] | 2.75 | 0.032 |
| stormal_median | [1.03, 5.54] | 2.39 | 0.042 | [1.57,28.47] | 6.69 | 0.01 |
| StrDiv | [1.02, 2.75] | 1.67 | 0.041 | [1.89, 9.52] | 4.24 | **<0.001** |
| TumCV | [0.27, 0.69] | 0.43 | **<0.001** | [0.10, 0.50] | 0.22 | **<0.001** |

In **Figure 2a**, we illustrated 3 important spatial features that can robustly stratify patients into statistically different prognostic groups, i.e., coefficient of variation of the tumor cell polygon area in the Voronoi diagram (TumCV), cellular diversity around stromal cells (StrDiv), and mean distance between stromal cells (StrDis). We demonstrated that, with the optimal cut-off values of these spatial features, 207 HCC patients from TCGA can be effectively stratified into better and worse prognostic groups (**Fig. 2b**). For example, for patients with TumCV lower than 35.8 (126 out of 207), the survival time longer than patients with TumCV higher than 35.8 (81 out of 207) in general. More importantly, with the same corresponding cut-off values, 67 patients from the independent Beijing Hospital cohort can also be effectively stratified into two prognostic groups (**Fig. 2c**). For example, for 207 HCC patients from TCGA, the HR between StrDiv - high group (StrDiv >0.11, 129 out of 207) and StrDiv -low group (StrDiv <0.11, 79 out of 207) is 1.67 (cox-p<0.05); for 67 HCC patients from Beijing Hospital, with the same cut-off value of StrDiv (0.11), the HR between StrDiv-high/low group (31 vs. 36) is 4.24 (cox-p<0.01). The same analysis method can be applied to StrDis (high group (129 out of 207) vs low group (78 out of 207) in TCGA cohort and high group (31 out of 67) vs low group (36 out of 67) in Beijing Hospital cohort). These results demonstrated that the developed spatial feature might robustly capture the prognosis-related spatial information inside the WSI across different cohorts. In summary, the spatial features developed in this study are effective and useful in patient stratification.

**Fig. 2.**
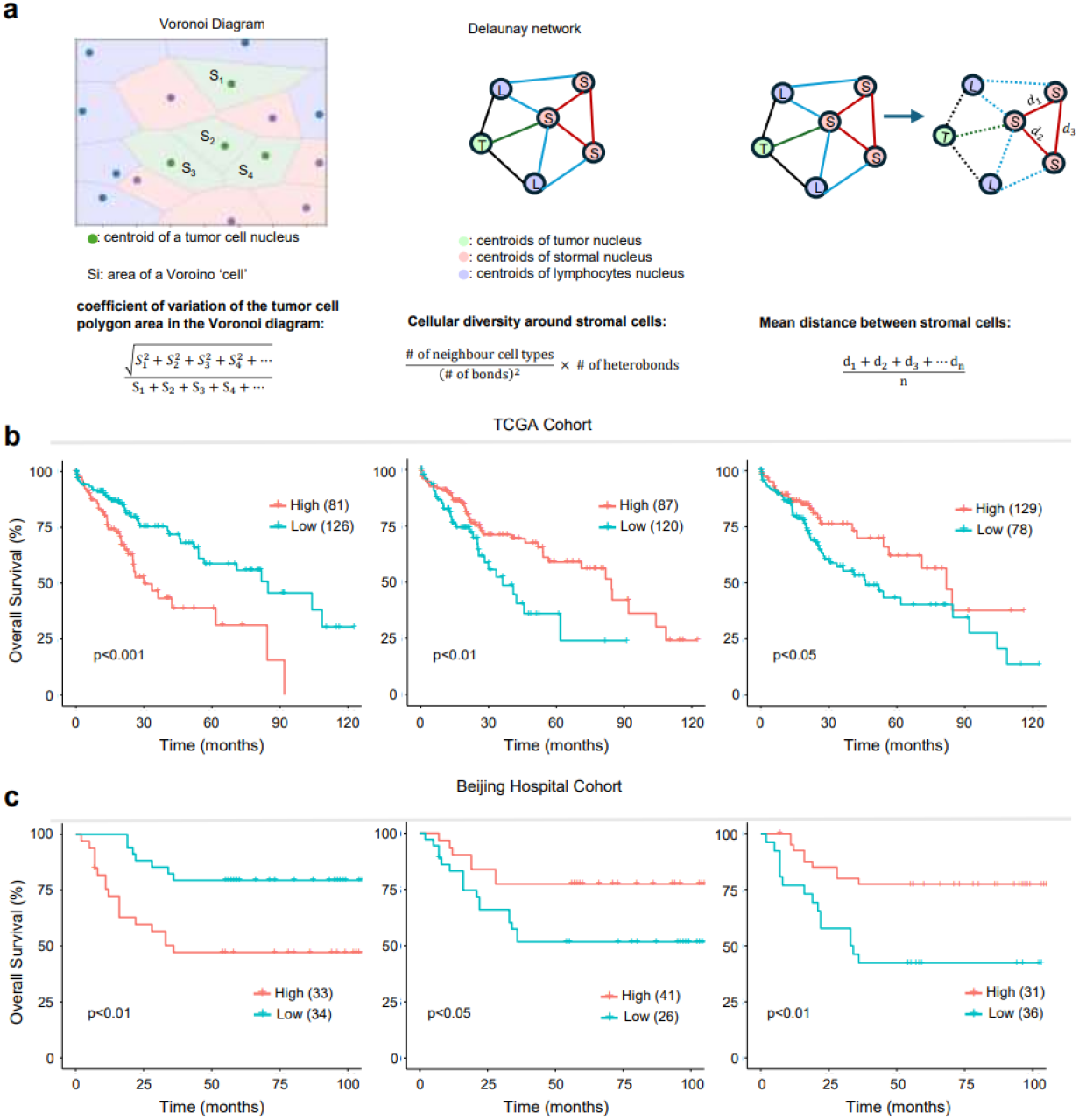
Analysis map of spatial features and Kaplan-Meier plots based on the same optimal cut-off grouping in TCGA dataset and Beijing Hospital dataset. (a) Illustration of features, left: coefficient of variation of the tumor cell polygon area in the Voronoi diagram (TumCV), middle: cellular diversity around stromal cells (StrDiv), right: mean distance between stromal cells (StrDis). (b)TCGA datasets, left: Kaplan-Meier plots based on TumCV, middle: Kaplan-Meier plots based on StrDiv, right: Kaplan-Meier plots based on StrDis. (c)Beijing hospital datasets, left: Kaplan-Meier plots based TumCV, middle: Kaplan-Meier plots based on StrDiv, right: Kaplan-Meier plots based on StrDis.

Moreover, so far, we simply stratified patients based on the median values of spatial topological features. included in the Cox model. To further test whether the features correlate with patient prognosis as continuum variables, we further performed Pearson correlation analysis using the TCGA-patients OS and the corresponding values of a specific feature (see **Methods**). As a result, ll_compass, ss_distance, StrDis, and stormal_density had a positive correlation coefficient with patient OS, with r values of 0.19, 0.16, 0.15, 0.14 respectively (all p<0.05). Conversely, OS was significantly negatively correlated with tumor_density, tumor_area, and TumCV with r values of -0.15, -0.16, -0.18 respectively, all P<0.05). These results also suggest that the developed spatial features in this work can be robustly associated with patient prognosis.

In addition, we also tested the consistency of patient stratification using clinic information, such as age, stage, *etc*. Note that TCGA and Beijing Hospital cohorts have different variables in two different cohorts. Only the TNM stage III correlated with patient prognosis on both cohorts (Supplementary Table S4). Therefore, according to the univariate analysis, our developed spatial features are effective and consistent for patient stratification.

### Multivariate analysis identifies TumCV and StrDiv as effective biomarkers

To systematically assess whether any of the spatial features could serve as the independent prognostic factors, we performed the multivariate analysis based on the spatial features listed in **Table 1** (13 of them) in addition to other clinical variables, such as gender, age, stage, treatment information, *etc*. A full list of corresponding variables and values was given in Supplementary Table S4.

The multivariate analysis for TCGA cohort showed that there are four spatial features having a p-value smaller than 0.1, with TumCV and StrDis having p-values smaller than 0.05. Note that even though the feature ll-distance also have a p-value smaller than 0.1, such feature also explicitly depends on the number of lymphocytes and therefore can suffer from the sampling fluctuations. Consequently, in the following analysis, we will only consider the top three independent spatial features identified by multivariate analysis. Next, we investigated whether a combination of these independent factors can enhance the efficacy of the patient stratification. Indeed, we demonstrated that patients with low TumCV and High StrDis tend to survive significantly longer than those with high TumCV and low StrDis (HR=3.4, p<0.001, Fig. 3b). Moreover, patients with low TumCV and high StrDiv tend to survive significantly longer than those with high TumCV and low StrDiv (HR=4.1, p<0.001, Fig. 3c). In addition, patients with High StrDis and high StrDiv tend to survive significantly longer than those with low StrDis and low StrDiv (HR=3.1, p<0.001, Fig. 3d). A combination of all 3 factors can further help to identify the worst (TumCV -high, StrDiv -low, StrDis -low) and best (TumCV -low, StrDiv -high, StrDis -high) prognostic groups (Supplementary Figure S1). These results demonstrate that the independent prognostic factors identified by the multivariate analysis can effectively enhance the patients stratification.

**Fig. 3.**
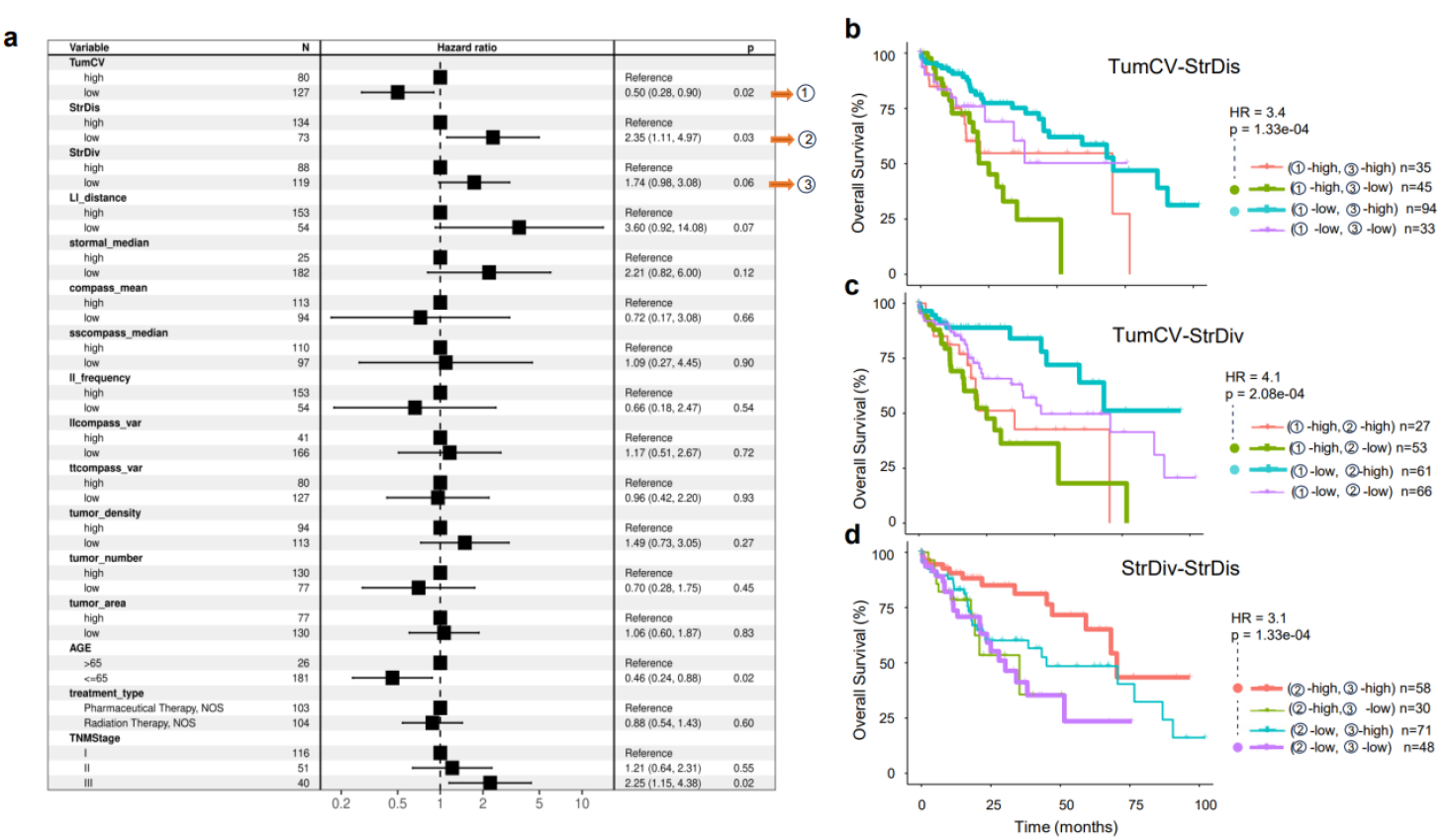
Multivariate Cox model and combine analysis of import features on TCGA cohort. (a), Forest plots of hazard ratios (estimates and 95 % CIs) from a multivariate Cox proportional hazards model of overall survival (OS) on TCGA cohort with both 13 significant factors and clinic variables about TCGA cohort. (b), Combine analysis of TumCV and StrDis. (c), Combine analysis of TumCV and StrDiv. (d), Combine analysis of StrDiv and StrDis.

Furthermore, we also performed multivariate analysis for the Beijing Hospital cohort. Based on the spatial features listed in Table 1, we first selected the ones that can also stratify patients from the Beijing Hospital cohort with the same cut-off values obtained from the univariate analysis of patients from TCGA cohort. In addition, we also include other clinical variables such as age, TNM Stage, MVI, HBV&HCV, AFP, *etc*. A full list of variables used for multivariate analysis is given in Supplementary Figure S2. The results indicates that there are two spatial features with their p-values smaller than 0.1, i.e., Tumor_density and StrDiv. Due to limited size of the Beijing Hospital cohort (67 patients), it is not surprising that the multivariate analysis results are not fully consistent with that of the TCGA cohort. However, it is worth noting that StrDiv is still a potential independent prognostic factor in the Beijing Hospital cohort, which further demonstrate the robustness of our developed spatial features in patient stratification across various cohorts. Furthermore, even though the p-values for TumVar is 0.29, it still ranks as the top 3 spatial features in the multivariate analysis. In addition to the spatial feature, results also show that no other clinical variable is an effective factor in predicting cancer prognosis.

### Impact of cellular topological features on the prognosis of patients with/without MVI

To further demonstrate the usefulness of spatial features in patient stratification, we investigated the effects of a combination of spatial features and known Clinical diagnostic criteria. Specifically, we chose MVI as an example to explore the potential improved efficacy. It has been demonstrated that MVI, which were classified based on whether microvascular invasion occurs [34], is an important prognostic index in clinic in HCC. However, for the 69-patient cohort from Beijing Hospital, MVI-positive/negative cannot effectively sperate patients into two prognostic groups with a logrank p value around 0.077 (**Fig. 4a**, upper panel). Therefore, we were curious whether the spatial features can help to understand why some of the patients without MVI had a worse prognosis.

**Fig. 4.**
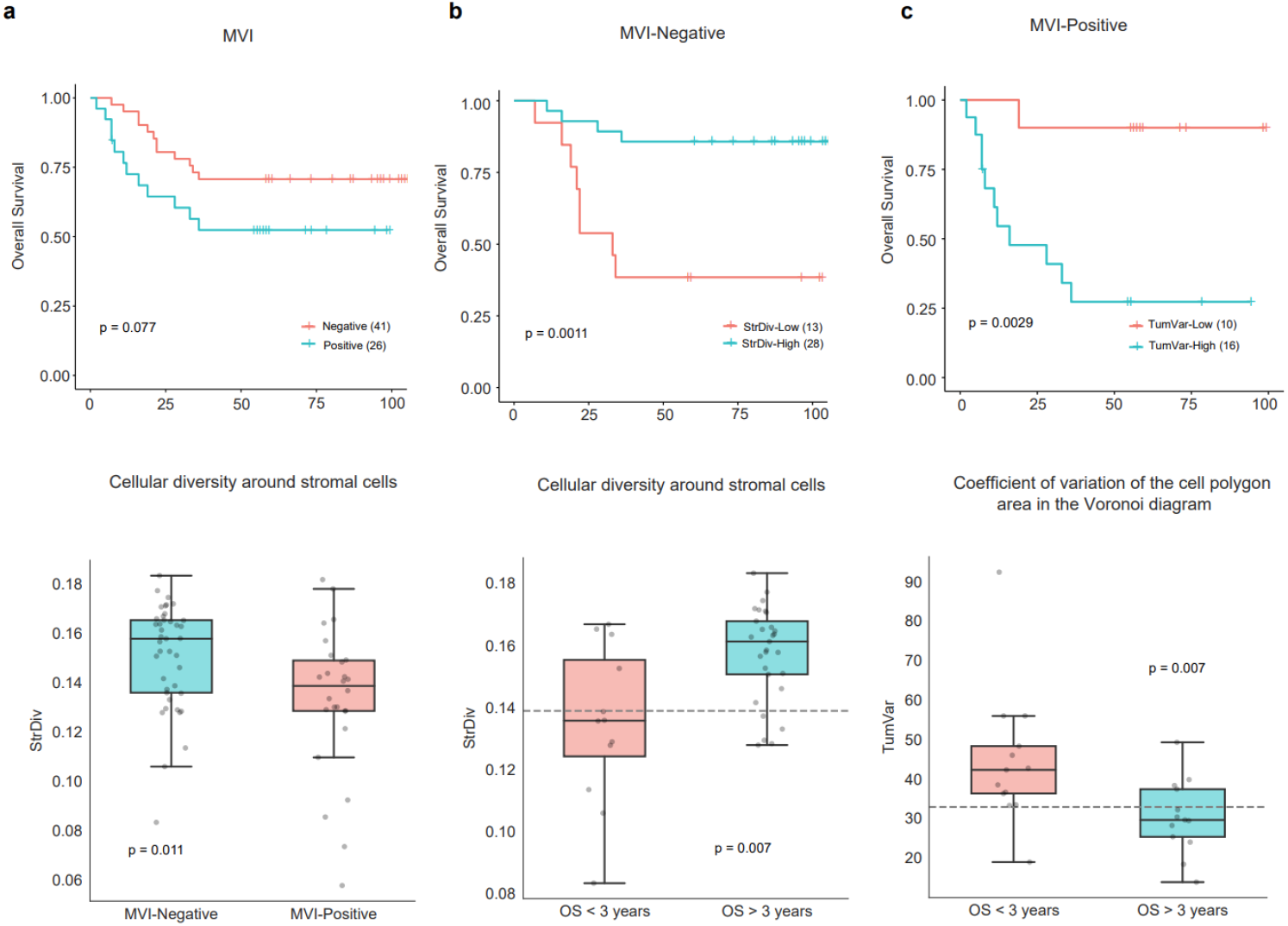
Impact of computed three features on the prognosis with MVI positive/ negative on Beijing hospital cohort. (a), Kaplan-Meier plots based on MVI subtype and the boxplot of StrDiv feature distribution grouped by MVI subtype. (b), Kaplan-Meier plots to show disease progression after dividing patients into high and low groups by StrDiv on MVI negative subtype and the boxplot of StrDiv distribution based on high/low survival subgroups by 36-months-cut-off on MVI negative subtype. (c), Kaplan-Meier plots to show disease progression after dividing patients into high and low groups by TumCV on MVI positive subtype and the boxplot of TumCV distribution based on high/low survival subgroups by 36-months-cut-off on MVI positive subtype.

We demonstrated that the cellular diversity around stromal cells (StrDiv), which is the overlapped spatial feature emerged from the multivariate analysis of both TCGA and Beijing-Hospital cohorts (p<0.1), has a statistical difference between MVI-positive/negative groups (**Fig. 4a**, bottom panel). For MVI positive patients, they tend to have a higher value of StrDiv. In **Figure 2a**, we have demonstrated that patients with a higher StrDiv tend to have a better prognosis. Therefore, such results suggest that our developed spatial features might have captured the prognosis-related information due to the existence of MVI. As controls, we also tested whether other clinical information could be different between the MVI positive/negative groups. But none of them showed a statistical difference.

To explore why some MVI-negative patients had a worse prognosis, we further applied the cut-off values of StrDiv obtained from TCGA cohort analysis to stratify the 46 MVI-negative patients into two groups and then performed the KM survival analysis. Interestingly, we showed that the spatial feature StrDiv can further stratify the MVI-negative patients, where the MVI-negative/StrDiv-high patients are more likely to have worse prognosis (**Fig. 4b**, upper panel). In addition, we also separated the MVI-negative patients into two groups with the OS longer or shorter than 3 years. The results in **Figure 4b** (lower panel) showed that there existed a statistically significant difference in the StrDiv values between the two patient groups. Therefore, these results demonstrated that combining the information of MVI and developed spatial features in this work, we can indeed further enhance the precision of patient stratification.

In addition to the enhanced stratification for the MVI-negative patients, we further explored whether the spatial features could help to better stratify MVI-positive ones. Specifically, we tested two spatial features, i.e., StrDiv and TumVar. Interestingly, we showed that patients with a high coefficient of variation of the tumor cell polygon area in the Voronoi diagram (TumCV) tend to have a significantly good prognosis among the MVI-positive patients (**Fig. 4c**, upper panel). Again, we also separated the MVI-positive patients into two groups with the OS longer or shorter than 3 years. Statistically, MVI-positive patients who survived longer than 3 years have a low TumCV, which is an indicator of better prognosis according to **Figure 2b**.

Therefore, these results demonstrated that combining the information of MVI and developed spatial features in this work, we can indeed further enhance the precision of patient stratification. Further test of the robustness of these findings could be performed in the future for an independent cohort with evaluated MVI information.

## Discussion

The tumor histology has also been demonstrated to be effective in prognostic stratification [5]. Recently, powered by deep-learning, image analysis of whole slide images (WSIs) of cancer tissue samples has become a new hotpot of biomedical research, where the characterization of the tumor microenvironment using the spatial information of cells inside the tumor can help better stratify patients into different prognostic groups, as well as predict their responses to therapies [35].

However, a notable limitation in previous studies is the lack of quantitative and explicit metrics to support the characterization of the tumor microenvironment. While the utilization of image analysis and deep learning has allowed for the extraction of valuable information from histological images, the absence of well-defined and quantifiable measures makes it challenging for clinicians to effectively interpret and apply the extracted image information. Consequently, there is a need for the development and implementation of robust and standardized metrics that can provide clear and measurable indicators of the tumor microenvironment’s characteristics.

In this work, we first tailored a previously reported CNN model to precisely obtain the locations of three types of cells in the H&E tumor-section images of HCC. Then by quantitatively defining the infiltration of TILs into the cancer-cell clusters, we showed that the characteristics of the structural heterogeneity and spatial organization changes of the tumor microenvironment were significantly correlated with the prognosis of patients.

Through the interpretation of clinical information features, we found that factors such as gender and staging are not effective predictors of patient survival. Additionally, treatment information such as chemotherapy or drug therapy does not show significant differences in patient survival. Therefore, it becomes particularly important to extract effective information from patient case slices and obtain additional key predictive factors for patient survival. In order to achieve quantitative cell-scale analysis in TIME (Tissue Image Analysis), we first establish a deep learning model to detect and identify different cell types, such as cancer cells, lymphocytes, and stromal cells. Then, we use topological graph analysis to quantify cell density, spatial distribution, and interactions based on the classified data. Finally, we use the calculated histological features to predict the clinical survival status of cancer patients.

The organization of the nucleus, such as the distribution, morphology, and arrangement of cells, has been shown to predict the invasiveness of cancer. For many different types of cancer, the hallmark of the disease is the disruption of structural cohesion between cancer cells and other primitive cells belonging to the same family. In contrast, more invasive tumors often exhibit lower levels of structural and organizational coherence between cells of the same primitive type compared to less invasive cancers. The features StrDiv and TumCV, constructed from our experimental results, have been validated.

In summary, in this work, by combining image analysis, deep-learning, and computer vision methodology, we developed a computational pipeline to use H&E pathological images to generate spatial-feature biomarkers to predict clinical outcomes, which also holds the promise for the application in other types of cancer as well as improving our understanding the cell-cell interactions in the tumor microenvironment. Our approach can provide crucial insights for designing novel cancer treatment methods. It can also expand our understanding of the impact of TME on the tumor evolution process. Our approach can be generalized to different cancer types, providing information for precision medicine strategies.

## Author contributions statement

H.H. performed the research and analyzed the results, X.L., J.S., and J.C. designed the research, T.T., J.C, and J.S. provided the clinical information of patients, H.H. and X.L. wrote the manuscript, and all authors reviewed and edited the manuscript.

## Acknowledgments

This work was supported by the National Key R&D Program of China (2021YFA0911100), the National Natural Science Foundation of China (32170672 and 32000886, 82073264), the Guangdong Basic and Applied Basic Research Foundation (2021A1515012461), and the Shenzhen Institute of Synthetic Biology (DWKF20210009).

