## Supplementary figures and images for "Deep learning-based Spatial Feature Extraction for Prognostic Prediction of Hepatocellular Carcinoma from Pathological Images"

### S2.png

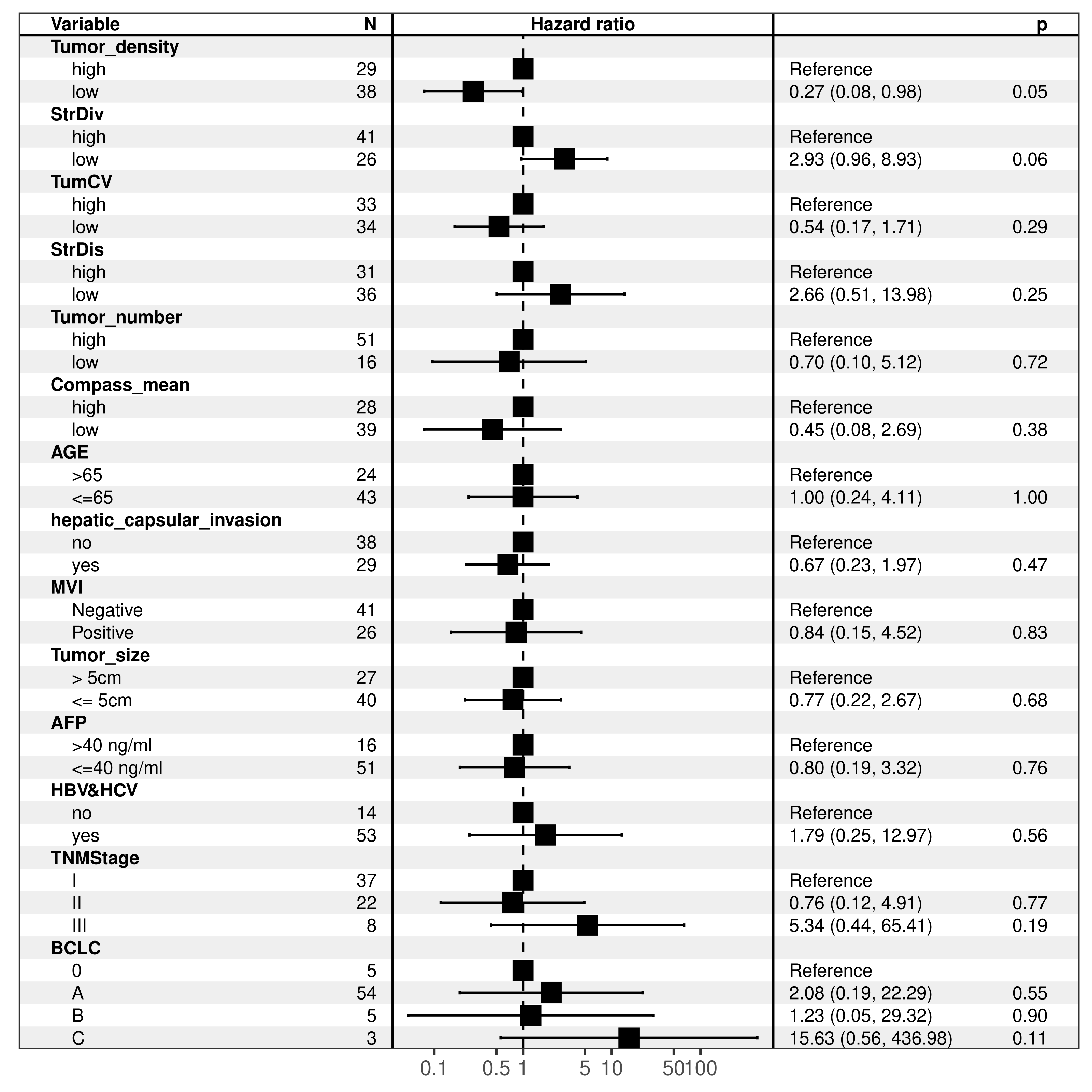
